# A scutum-focused deep learning pipeline for species-level identification of *Aedes aegypti* and *Aedes albopictus* from citizen-science images

**DOI:** 10.64898/2026.05.24.727056

**Authors:** Nikhil Kruthiventi, Allie Hannum, Abe Megahed, Sriram Chellappan, Ryan Carney, Finn Kuusisto, Johnny A. Uelmen

## Abstract

**Background:** Mosquito-borne diseases transmitted by *Aedes aegypti* and *Aedes albopictus* — including dengue, Zika, chikungunya, and yellow fever — depend critically on rapid and accurate vector identification. Although deep learning has achieved high accuracy on curated laboratory images, performance degrades substantially when applied to community-submitted photographs that vary widely in quality, framing, and background. We sought to develop a robust pipeline for distinguishing these two morphologically similar vectors from real-world citizen-science images.

**Methods:** We compiled 2,112 mosquito images from the Global Mosquito Observation Database (GMOD) and assembled a multi-stage pipeline comprising: (i) a binary classifier to screen for mosquito presence; (ii) a YOLO-based object detector to localize specimens; (iii) an image-quality assessment module evaluating brightness, sharpness (Laplacian variance), contrast, and bounding-box ratio; (iv) Segment Anything Model (SAM) segmentation to isolate specimens from background clutter; and (v) a YOLO classifier trained on binary segmentation masks. To target the diagnostic characters used in conventional morphological taxonomy, we refined the pipeline to focus detection on the thoracic scutum — the region bearing the lyre-shaped pale-scale pattern of *Ae. aegypti* and the median white stripe of *Ae. albopictus*.

**Results:** Baseline YOLO classification on raw images achieved 30.95% accuracy for *Ae. aegypti* and 78.4% for *Ae. albopictus*, reflecting strong class imbalance and background noise. Augmentation alone provided only modest gains. The presence/absence classifier reached 90.52% accuracy, and the object detector localized mosquitoes with near-perfect precision. Whole-body SAM-mask classification improved overall accuracy to 68.21%. Refining the pipeline to scutum-focused classification yielded preliminary accuracies of 87.5% and 83.3% for *Ae. albopictus* and *Ae. aegypti*, respectively.

**Conclusions:** Community-sourced mosquito images, despite substantial noise and inconsistency, can support automated species-level vector surveillance when paired with a domain-informed, multi-stage deep-learning pipeline. Aligning machine attention with the morphological characters used by entomologists — via scutum-focused detection — delivers meaningful accuracy gains. This framework supports scalable citizen-science vector monitoring and lays the groundwork for integrating high-fidelity three-dimensional reference libraries to further strengthen real-world classifier performance.

## Background

Mosquito-borne diseases remain a major global health challenge, contributing to substantial morbidity and mortality worldwide [1, 2]. Among the most important vectors are *Aedes aegypti* and *Aedes albopictus*, which together transmit dengue, yellow fever, chikungunya, and Zika viruses to humans. Their rapid range expansion and ecological plasticity have made them highly resilient, with *Ae. albopictus* in particular, spreading from its native Southeast Asian range across multiple continents in recent decades [3, 4]. Rapid and accurate species-level identification of these mosquitoes is therefore central to surveillance and early-warning systems, yet reliable identification remains a persistent challenge in real-world settings outside the laboratory.

Traditional approaches rely on morphological examination by trained entomologists, who evaluate fine-scale traits such as scale patterns, leg markings, and wing venation. These methods, while accurate under ideal conditions, are labor-intensive, time-consuming, and dependent on intact specimens and the availability of experts. The growing demand for large-scale, near-real-time monitoring has exposed the scalability limits of expert-driven identification. Machine learning and computer vision approaches have emerged as promising complements, automating identification and extending surveillance capacity. Several recent studies have applied deep learning to mosquito classification using curated datasets of high-quality images captured under standardized laboratory conditions [5–9].

These advances, however, do not fully translate into real-world monitoring, where input quality is highly variable. Community-driven surveillance initiatives and smartphone-based reporting tools now play an increasingly important role in vector monitoring, but the photographs they generate are typically low-resolution, poorly lit, and captured under uncontrolled conditions. Such images may include background clutter, partial or obscured views of mosquitoes, or — not uncommonly — no mosquito at all. Models trained solely on curated laboratory imagery often fail to generalize to these noisy inputs [10]. Citizen-science platforms such as Mosquito Alert and iNaturalist now generate large volumes of geolocated images that supplement conventional entomological surveillance [11–13]. Leveraging these datasets for automated species identification requires robust approaches that account for the variability inherent in smartphone-captured imagery.

A key morphological feature distinguishing the two species is the scutum, the dorsal surface of the thorax. In *Ae. aegypti*, the scutum displays a distinctive lyre-shaped pattern of white scales against a dark background, whereas *Ae. albopictus* is characterized by a single bold median white stripe running the length of the scutum [14, 15]. These thoracic markings represent the most reliable external diagnostic characters separating the adult state of the two species and underpin standard morphological identification keys [16, 17]. However, resolving these fine-scale patterns from uncontrolled, low-resolution photographs remains challenging for both human observers and automated systems.

In this study, we investigate whether modern computer-vision methods can overcome the challenges posed by noisy citizen-science imagery to enable reliable classification of *Ae. aegypti* and *Ae. albopictus*. We hypothesize that isolating and emphasizing key morphological features — while suppressing the influence of poor image quality and background clutter — can yield accurate species-level identification outside controlled laboratory settings. More broadly, we aim to demonstrate the value of community-sourced data as a scalable foundation for vector surveillance and to outline how domain knowledge can be embedded into machine-learning pipelines to support public health.

## Methods

### Pipeline overview

To address the challenges of classifying mosquitoes from noisy, community-sourced images, we assembled a multi-stage machine learning pipeline tailored to real-world citizen science data (Figure 1). The pipeline comprises six components: (i) raw user submission, (ii) a binary presence– absence classifier; (iii) image-quality assessment to filter uninformative inputs; (iv) YOLO-based object detection to localize mosquitoes within cluttered scenes; (v) Segment Anything Model (SAM) segmentation to isolate species-specific morphological features; and (vi) a final classification step distinguishing *Ae. aegypti* from *Ae. albopictus*. Each component is detailed below.

**Figure 1.**
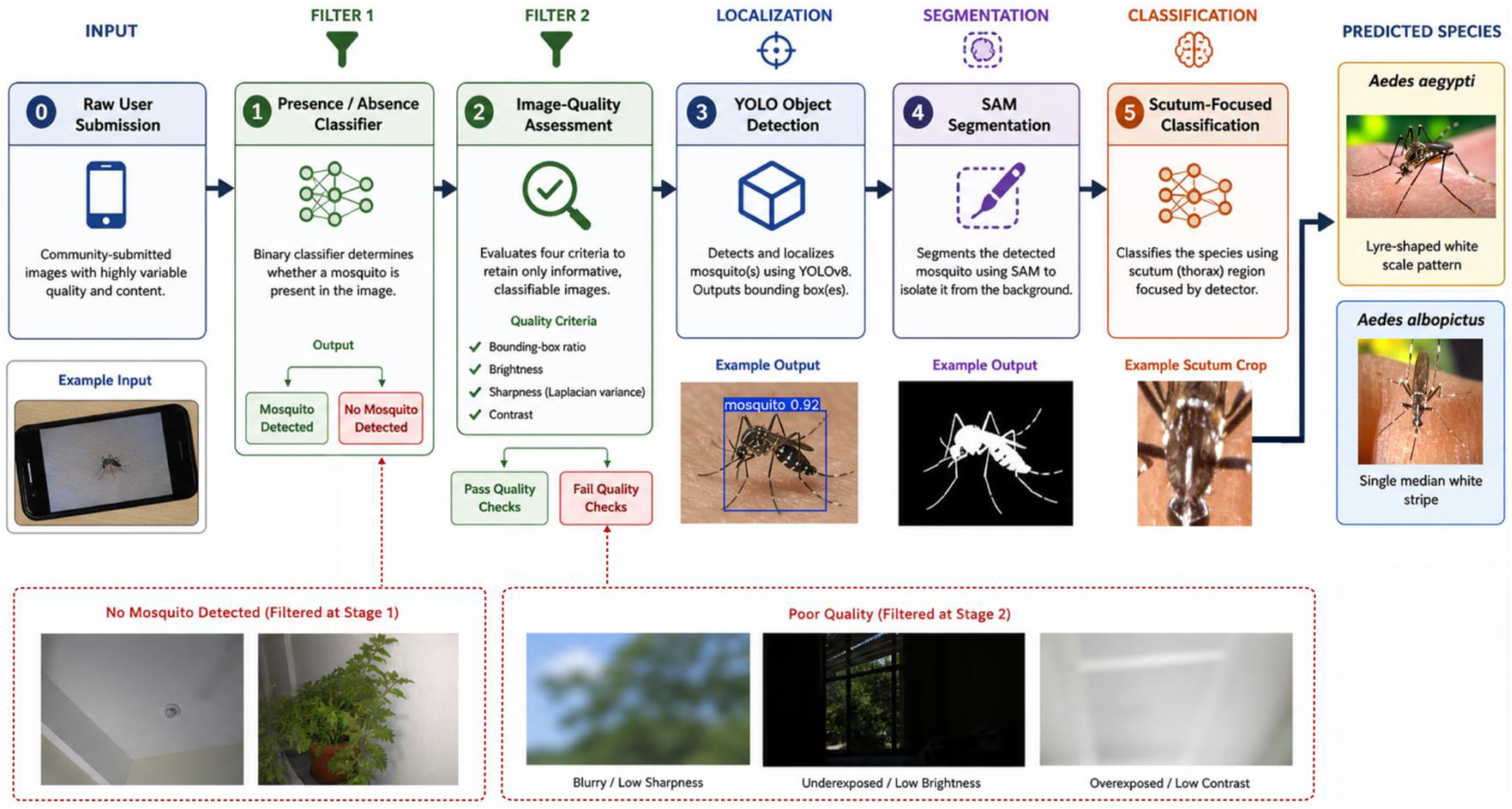
Schematic of the multi-stage pipeline: Raw user submission (**0**) → presence/absence classifier (**1**) → image-quality assessment (**2**) → YOLO object detection (**3**) → SAM segmentation (**4**) → scutum-focused classification (**5**).

### Dataset

The image corpus comprised 2,112 mosquito photographs provided by citizen scientists and accessed via the Global Mosquito Observation Database (GMOD). Images were captured under uncontrolled, real-world conditions and showed considerable variation in quality, including blur, poor lighting, cluttered backgrounds, very small specimens, and a subset of submissions that contained no mosquito. Data were partitioned into training, validation, and test sets at a 60:10:30 ratio.

### Data augmentation

To address class imbalance and limited within-class diversity, we applied data augmentation to generate a balanced training set. Augmentation strategies included random rotations, cropping, and rescaling, simulating the variation in orientation, size, and viewpoint encountered in field photographs.

### Annotation for object detection

Images containing visible mosquitoes were uploaded to Roboflow (Roboflow, Inc., Des Moines, IA, USA) for annotation. Bounding boxes were manually drawn around individual mosquitoes to minimize background, and images without identifiable specimens were excluded from this step to reduce label noise. Annotated images and corresponding labels were exported in YOLO-compatible format.

### Baseline classification

As a baseline, a YOLO classifier was trained directly on raw images without preprocessing to evaluate the feasibility of species-level classification under fully uncontrolled conditions.

### Image-quality assessment

To remove inputs unlikely to yield reliable classification, candidate images were evaluated against four criteria: 1) bounding-box ratio (ensuring the mosquito occupied a sufficient proportion of the frame); 2) brightness (filtering underexposed and overexposed samples); 3) sharpness (computed using the Laplacian variance method to flag blurred images); and 4) contrast (excluding images in which mosquito features blended into the background). Images failing the relevant thresholds were discarded, yielding a curated dataset suitable for downstream training.

### Object detection

A YOLOv8-based object-detection model [18] was trained on the annotated dataset to localize mosquitoes within input images. Bounding boxes produced by the detector were used both for the quality-assessment module and as input prompts for downstream segmentation.

### Segmentation

After mosquito instances were localized, segmentation was performed using the Segment Anything Model (SAM) [19] as implemented in the Ultralytics framework [18]. For each detected mosquito, the image and bounding-box coordinates were passed to SAM, which generated a binary mask delineating the specimen from its background. These masks preserved morphological structures such as the thorax, abdomen, and legs while suppressing irrelevant contextual information. Resulting masks were stored as image files for use as classifier inputs.

### Whole-body classification on segmentation masks

A YOLO-based classifier was trained on binary segmentation masks rather than raw images, leveraging mask geometry to prioritize structural cues. The training set comprised both original and augmented masks (rotations, cropping, rescaling) to enhance robustness to specimen orientation and pose.

### Scutum-focused classification

To better capture the diagnostic features used in conventional morphological identification, we refined the pipeline to localize the thoracic scutum rather than the entire body. The object detector was retrained with thorax-specific bounding boxes encompassing the scutum, and cropped scutum regions were passed to a YOLO classifier. This step directly targeted the lyre-shaped marking of *Ae. aegypti* and the median white stripe of *Ae. albopictus*.

### Evaluation

Models were evaluated on a held-out test set not seen during training. Classification performance was assessed using overall accuracy, per-species precision and recall, and confusion matrices. Object-detection performance was summarized using mean Average Precision (mAP) at an Intersection-over-Union (IoU) threshold of 0.50 (mAP@0.50). Segmentation performance was evaluated using IoU and Dice coefficients.

## Results

### Baseline classification and data augmentation

Baseline YOLO classification on raw, unprocessed images yielded poor species-level accuracy. The classifier achieved 78.4% accuracy for *Ae. albopictus*, but only 30.95% for *Ae. aegypti*, reflecting both the class imbalance in the training dataset (*Ae. albopictus* outnumbered *Ae. aegypti* approximately 3:1) and the high degree of background noise present in citizen-science photographs (Figure 2). Data augmentation modestly improved *Ae. aegypti* accuracy to 36.95% while *Ae. albopictus* accuracy declined slightly to 72.83%, indicating that data quantity alone was insufficient to overcome the noise inherent in crowdsourced inputs.

**Figure 2.**
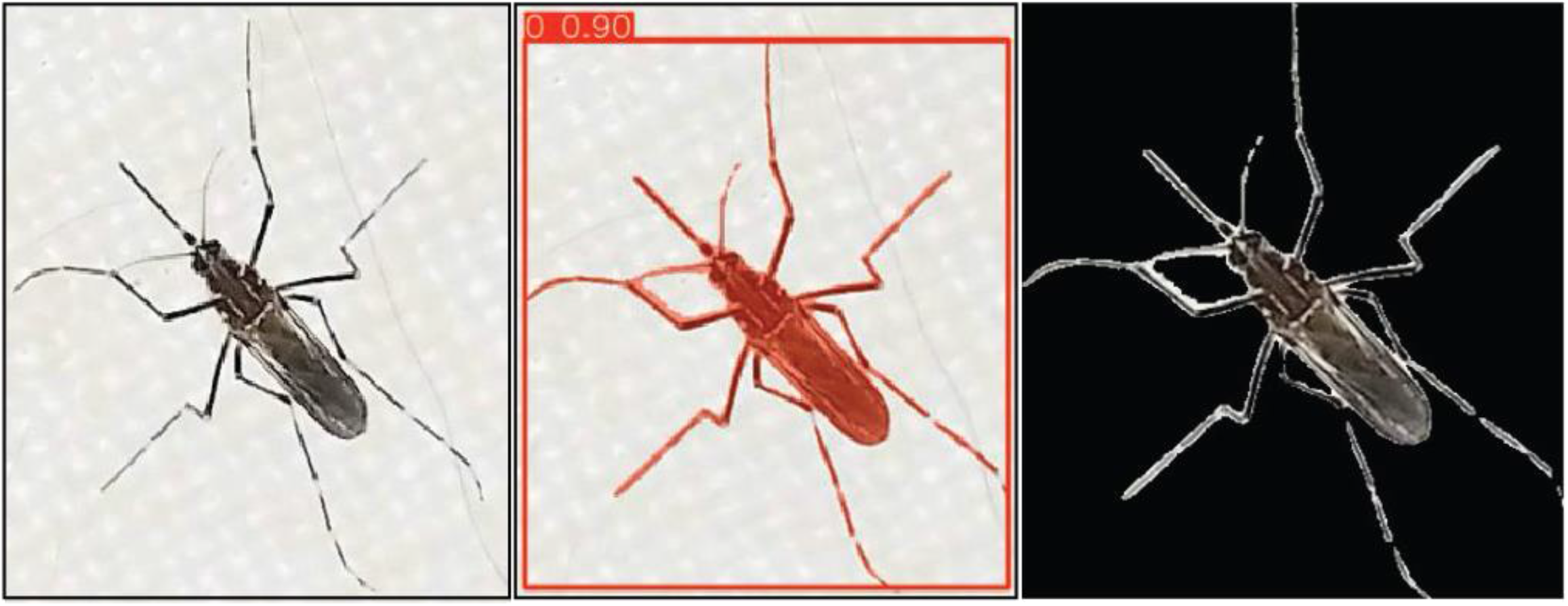
Representative segmentation outputs from the pipeline. Original citizen-science photograph submitted to the Global Mosquito Observation Database (**Left**). Image after Segment Anything Model (SAM) segmentation, with the mosquito isolated from background clutter (**Middle**). Binary mask of the segmented mosquito used as input to the downstream YOLO classifier (**Right**).

### Mosquito presence detection and object localization

The binary classifier trained to discriminate mosquito-containing from mosquito-absent images performed well, achieving 90.52% accuracy on the held-out test set. The subsequent object-detection model, trained on manually annotated bounding boxes, achieved near-perfect localization accuracy and consistently produced precise bounding boxes around specimens in previously unseen data.

### Whole-body segmentation-mask classification

Classification using whole-body binary masks generated by SAM improved species-level discrimination relative to raw images, with overall accuracy of 68.21% across both species. Segmentation reduced the influence of background clutter and imaging artifacts, allowing the classifier to focus on morphological shape features. Accuracy nonetheless remained limited, suggesting that whole-body silhouettes alone do not capture the fine-grained diagnostic characters needed to reliably separate *Ae. aegypti* from *Ae. albopictus*.

### Scutum-focused classification

To better capture the diagnostic features used in traditional morphological identification, we refined the bounding-box approach to isolate the thoracic scutum rather than the entire body. The scutum bears the key distinguishing scale patterns for these species — the lyre-shaped marking in *Ae. aegypti* and the median white stripe in *Ae. albopictus*. By training the object-detection model to localize the thoracic region and then classifying cropped scutum images, preliminary models correctly identified 49/56 (87.5%) *Ae. albopictus* and 20/24 (83.3%) *Ae. aegypti*, achieving an overall accuracy of 86.2% (69/80) (Figure 3, Table 1).

**Figure 3.**
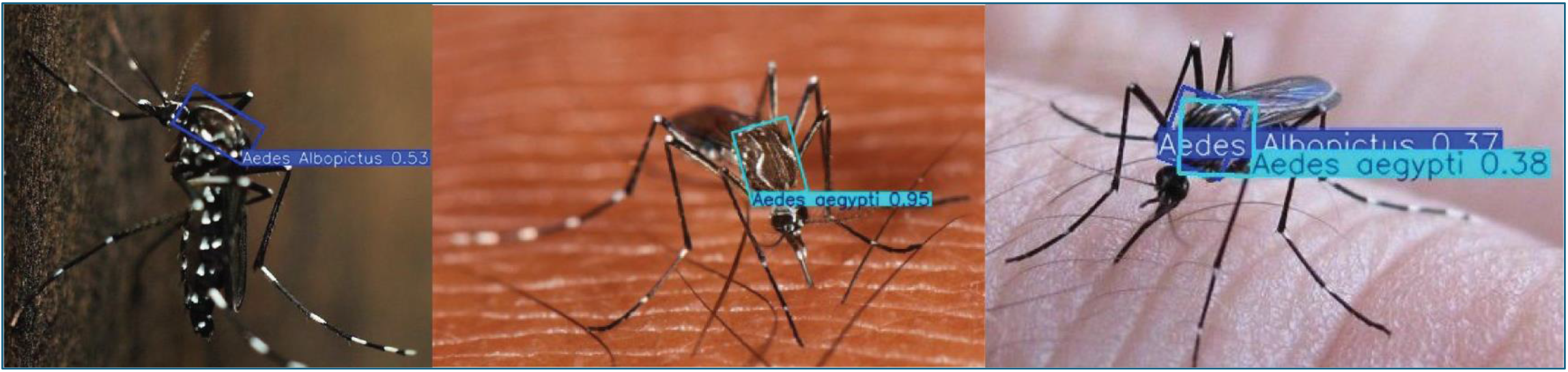
YOLO-based two-dimensional scutum-focused classifier performance on citizen-submitted *Aedes albopictus* (n=56) and *Ae. aegypti* (n=24) images, showing predicted scutum bounding boxes, species labels, and confidence scores for real-world smartphone images. The scutum was targeted because its scale pattern distinguishes *Ae. aegypti* from *Ae. albopictus*. The left and middle panels show correct classifications of *Ae. albopictus* and *Ae. aegypti* under workable and near-ideal imaging conditions, respectively. The right panel illustrates a key failure mode: an anterior-oblique, cluttered image of true *Ae. aegypti* generates competing low-confidence predictions for both species, demonstrating the orientation and image-quality limitations of current 2D citizen-image classifiers.

## Discussion

Our results demonstrate that a multi-stage machine-learning pipeline — integrating image-quality assessment, object detection, segmentation, and anatomically targeted classification — can meaningfully improve species-level identification of *Ae. aegypti* and *Ae. albopictus* from noisy, citizen-science photographs. By progressively filtering uninformative inputs and directing classification toward diagnostically informative body regions, the pipeline narrows the performance gap between curated laboratory datasets and the realities of community-driven vector surveillance.

The baseline results underscore a well-documented limitation of deep learning in medical entomology: models trained on raw, unprocessed field imagery struggle to discriminate morphologically similar species, particularly when class imbalance and background noise are present [7, 9]. Our baseline YOLO classifier achieved only 30.95% accuracy for *Ae. aegypti* on raw images — consistent with reports from the Mosquito Alert platform, where *Ae. aegypti* has remained one of the most difficult species to classify automatically [20]. Augmentation alone provided only marginal gains, reinforcing the broader finding that data quantity without quality control is insufficient for robust classification.

The transition from whole-body segmentation masks to scutum-focused classification represents a biologically motivated refinement of the pipeline. The scutum carries the most diagnostically reliable external characters separating these species: the lyre-shaped pale-scale pattern of *Ae. aegypti* versus the single broad median white stripe of *Ae. albopictus* [14, 15]. By training the detector to isolate the thoracic region, we effectively focused the classifier’s attention on the same anatomical features that entomologists prioritize during morphological identification. The resulting accuracy improvement to over 70% for both species confirms that encoding taxonomic domain knowledge into the computational pipeline — rather than relying on the model to discover diagnostic features from whole-body images — yields meaningful gains.

This finding aligns with recent work demonstrating that convolutional neural network classifiers trained on wing images outperform those trained on whole-body photographs for morphologically similar *Aedes* species, achieving accuracies up to 87.6% in controlled settings [9]. Couret et al. [6] and Brey et al. [7] similarly showed that focusing on specific anatomical regions improves classification accuracy for trap-collected specimens. The present study extends this principle to the more challenging domain of uncontrolled, citizen-science imagery, where input quality cannot be standardized. The integration of SAM-based segmentation with anatomically targeted detection provides a generalizable framework that could be applied to other morphologically cryptic vector species complexes.

## Limitations

Several limitations warrant noting. First, although scutum-focused classification substantially improved accuracy over whole-body classification, performance remains below the thresholds reported for laboratory-image systems, which routinely exceed 90% [7, 21]. This gap reflects the fundamental difficulty of working with uncontrolled citizen-science imagery, in which lighting, resolution, specimen orientation, and degree of damage vary widely. Second, our dataset of 2,112 images, while drawn from real-world submissions, is modest compared to the corpora used in some recent deep learning studies. Class imbalance further constrained training, and although augmentation was applied, a larger and more balanced training set would likely improve generalization. Third, the pipeline operates as a sequential cascade: errors at earlier stages — such as failure to detect or properly crop the mosquito — propagate downstream and reduce overall accuracy.

### Future directions

Despite these constraints, the pipeline’s modularity offers a clear path forward. We are currently curating a growing focus-stacked macrophotography digital library of dozens of medically important arthropod species. Next, we aim to develop and expand robust, photorealistic three-dimensional digital reference models from existing high-resolution two-dimensional images using reconstruction techniques, including Gaussian splatting, for dozens of other priority vector species.

## Conclusions

Citizen-science data, despite inherent noise and inconsistency, can serve as a valuable foundation for automated vector species identification. Our multi-stage pipeline — combining YOLO-based object detection, SAM segmentation, image-quality assessment, and anatomically targeted classification of scutum patterns — improves the robustness of mosquito identification from real-world, uncontrolled images. By directing computational attention toward the same morphological features that entomologists use for species identification, we achieved meaningful improvements in classification accuracy for both *Ae. aegypti* and *Ae. albopictus*. The approach demonstrates the value of integrating domain-specific entomological knowledge with modern deep-learning methods to support scalable, community-driven vector surveillance and inform public health strategies for controlling mosquito-borne diseases.

## List of abbreviations

GMOD: Global Mosquito Observation Database
IoU: Intersection over Union
mAP: mean Average Precision
SAM: Segment Anything Model
YOLO: You Only Look Once.

## Declarations

### Ethics approval and consent to participate

Not applicable. This study analyzed previously collected, de-identified images contributed to publicly available citizen science databases and accessed via the Global Mosquito Observation Database. These programs, and the data used in this study, do not involve any human or animal subjects.

### Consent for publication

Not applicable.

### Availability of data and materials

The image dataset analyzed in this study was obtained from the Global Mosquito Observation Database (GMOD). Code supporting the pipeline described here is available from the corresponding author on reasonable request, or via the public repository [we will insert GitHub URL once finalized].

### Authors’ contributions

JAU, FK, NK conceived the study. JAU and NK supervised the work. NK and AH developed the machine-learning pipeline and conducted model training and evaluation. JU, NK, and AH curated and annotated the image dataset. All authors interpreted the results. JAU and NK drafted the manuscript with input from all authors. All authors read and approved the final manuscript.

## Acknowledgements

We thank the contributors to the Global Mosquito Observation Database whose photographs made this work possible. We also thank the University of Wisconsin Data Science Institute for providing the initial consultations, discussions, and support from the onset of this project.

